# SARS-CoV-2 spike protein induces endothelial dysfunction in 3D engineered vascular networks

**DOI:** 10.1101/2022.10.01.510442

**Authors:** Brett Stern, Peter Monteleone, Janet Zoldan

**Affiliations:** The University of Texas at Austin, Department of Biomedical Engineering, Austin, Texas; The University of Texas at Austin, Dell Medical School, Department of Internal Medicine, Austin, Texas; Ascension Texas Cardiovascular, Austin, Texas

## Abstract

With new daily discoveries about the long-term impacts of COVID-19 there is a clear need to develop *in vitro* models that can be used to better understand the pathogenicity and impact of COVID-19. Here we demonstrate the utility of developing a model of endothelial dysfunction that utilizes induced pluripotent stem cell-derived endothelial progenitors encapsulated in collagen hydrogels to study the effects of COVID-19 on the endothelium. We found that treating these cell-laden hydrogels with SARS-CoV-2 spike protein resulted in a significant decrease in the number of vessel-forming cells as well as vessel network connectivity. Following treatment with the anti-inflammatory drug dexamethasone, we were able to prevent SARS-CoV-2 spike protein-induced endothelial dysfunction. In addition, we confirmed release of inflammatory cytokines associated with the COVID-19 cytokine storm. In conclusion, we have demonstrated that even in the absence of immune cells, we are able to use this 3D *in vitro* model for angiogenesis to reproduce COVID-19 induced endothelial dysfunction seen in clinical settings.

## Introduction

The ongoing COVID-19 pandemic, caused by the novel coronavirus SARS-CoV-2, has as of the time of writing infected 605 million people, 6.5 million of whom have died^1^. Although initial focus was placed on the virus’ effects on the lungs, it is now known to affect the endothelium as well, especially in severe infection^2^. Severe COVID-19 is associated with varied vascular pathologies, including microthrombi, complement pathway activation, and endothelial cell sloughing from the vessel wall. This is known as endothelial dysfunction, and presence of endothelial dysfunction is associated with a 5-10-fold increase in mortality rate^3^. Additionally, these mechanisms have led to profound vascular and cardiac morbidity secondary to COVID-19^4,5^. COVID-19 mediated endothelial dysfunction occurs as a result of SARS-CoV-2’s binding to the ACE2, a surface protein involved in blood pressure regulation, resulting in downstream signaling that promotes inflammation^2^. This is in contrast to other coronaviruses, since although they are able to infect endothelial cells and promote an inflammatory environment, extensive vascular damage is unique to SARS-CoV-2^6^. To better understand the interactions between SARS-CoV-2 and endothelial cells and to evaluate the effectiveness of proposed treatments, there is a need to create *in vitro* tissue models of SARS-CoV-2-induced endothelial dysfunction.

Other research groups have created *in vitro* tissue models for SARS-CoV-2’s effects on the endothelium^3,7^. Following exposure to the virus, researchers have observed viral infection of endothelial cells, decreases in expression of cell adhesion molecules leading to disruption of the endothelial barrier, and increased secretion of inflammatory cytokines. However, existing models for the effects of SARS-CoV-2 on the endothelium were developed using cells cultured in 2D monolayers, which do not model conditions *in vivo* as accurately compared to 3D models; culturing cells in 2D rather than 3D results in significant changes in proliferation rate, cell morphology, gene expression, and drug responsiveness^8,9^. In addition, endothelial cells cultured in 2D cannot form vascular networks, which is a key feature for identifying healthy and functional cells; this is especially important for COVID-19 modelling, where vascular network disruption is a hallmark of viral infection^6^.

Previously, our group has utilized human induced pluripotent stem cells (hiPSCs) differentiated to endothelial progenitors (EPs) and encapsulated in collagen hydrogels to study angiogenesis^10-12^. Here we utilize this model to demonstrate that hiPSC-EP laden hydrogels treated with SARS-CoV-2 spike protein (CSP) results in significant vascular disruption, both in terms of the number of cells part of the vascular networks as well as vascular connectivity. We also measured increases in inflammatory cytokine release consistent with that observed in clinical settings as well as a reduction in CSP-induced vascular dysfunction following treatment with the anti-inflammatory drug dexamethasone. In conclusion, we are able to reproduce COVID-19-induced endothelial dysfunction seen in clinical settings using hiPSC-EP-laden hydrogels treated with CSP.

## Materials and Methods

### Maintenance of hiPSCs

Human indued pluripotent stem cells (hiPSCs), derived from human dermal foreskin fibroblasts (DF19-19-9-11T), were purchased from WiCell. iPSCs were cultured under feeder-free conditions on vitronectin-coated (ThermoFisher Scientific) six-well plates in complete Essential 8 (E8) medium (ThermoFisher). hiPSCs were passaged upon reaching 70–80% confluency. To passage, hiPSCs were rinsed with Dulbecco’s phosphate-buffered saline (DPBS) and then treated with 0.5 mM ethylenediaminetetraacetic acid (EDTA) for 4.5 minutes at 37°C. EDTA was then removed and the hiPSCs were resuspended in complete E8 and seeded as small colonies onto freshly coated vitronectin plates.

### Differentiation of hiPSCs to CD34^+^ hiPSC-EPs

hiPSCs were differentiated into hiPSC-EPs following an established protocol from Lian *et al* with small modifications^13^. In brief, iPSCs were manually dissociated into a single-cell suspension in complete E8 supplemented with 10 μM ROCK Inhibitor (Y-27632; Selleckchem) and plated on Matrigel-coated 24-well plates at a density of 20,000 cells/cm^2^. 24 hours after seeding, media was replaced with complete E8 without ROCK Inhibitor. 48 hours after seeding, media was replaced with LaSR Basal (Advanced Dulbecco’s Modified Eagle Medium (DMEM)/F12 (ThermoFisher Scientific) supplemented with 60 μg/mL L-ascorbic acid 2-phosphate sesquimagnesium salt (Sigma-Aldrich) and 2.5 mM GlutaMAX (ThermoFisher Scientific)) with 6 μM CHIR99021 (LC Technologies). LaSR Basal plus CHIR99021 was replaced after 24 hours. 48 hours after CHIR99021 induction, media was replaced with fresh LaSR Basal without CHIR99021 for three additional days.

### Fluorescence-activated cell sorting of CD34-expressing hiPSC-EPs (CD34^+^ hiPSC-EPs)

Five days after CHIR99021 induction, the differentiated cells were incubated with Accutase (ThermoFisher Scientific) for 10 minutes at 37°C and manually dissociated into single cells. Cells were centrifuged at 300g for 5 minutes and the pellet was resuspended in 200 μL of sorting buffer (2 mM EDTA and 0.5% bovine serum albumin (BSA; Sigma-Aldrich) in DPBS) and 2 μL of CD34-PE antibody (Miltenyi Biotec). Cells were then incubated at 4°C for 10 minutes. Cells were filtered twice with a 35 μm cell strainer (Corning) to remove cell and extracellular matrix aggregates, and CD34^+^ hiPSC-EPs were isolated using a fluorescence-activated cell sorting (FACS) instrument (S3e; Bio-Rad). Gating and population analysis were performed with Bio-Rad software native to the S3e cell sorter.

### Encapsulation of CD34^+^ hiPSC-EPs in collagen hydrogels

A total of 500,000 CD34^+^ hiPSC-EPs were centrifuged twice and resuspended in Complete Endothelial Growth Media 2 (EGM-2, Lonza) supplemented with 50 ng/mL recombinant human vascular endothelial growth factor (VEGF; R&D Systems), 10 μM Y-27632, and 100 U/mL penicillin-streptomycin (ThermoFisher Scientific), termed encapsulation media. Ice-cold 10 mg/mL collagen (Type I, rat tail; Corning), 10x Medium 199 (ThermoFisher Scientific), and additional encapsulation media were mixed together, and the collagen was neutralized with 1M sodium hydroxide (NaOH; Sigma-Aldrich), turning the solution bright pink. The CD34^+^ hiPSC-EP solution was then added to the collagen solution. The final concentrations for all collagen hydrogel components were described previously^10-12^. 56 μL of the collagen-cell suspension was pipetted into each well of an ultra-low attachment round-bottomed 96-well plate (Nexcelom) and allowed to solidify for 30 minutes at 37°C and 5% CO2. CD34^+^ hiPSC-EP-laden hydrogels were immersed in 100 μL of encapsulation media. 24 hours after cell encapsulation, media was replaced with complete EGM-2 supplemented with 50 ng/mL VEGF, termed endothelial culture media, which was replaced daily for 7 days.

### Immunocytochemistry

The vessel-like networks generated within CD34^+^ iPSC-EP-laden hydrogels were visualized by following an established immunocytochemistry protocol for cells embedded in thick collagen hydrogels^10-12^. In brief, CD34^+^ iPSC-EP-laden hydrogels were fixed in a 4% paraformaldehyde (PFA; Polysciences) for 10 minutes, washed three times with DPBS, and then permeabilized in 0.2% Triton X-100 (Amresco) for 10 minutes. After three washes with DPBS supplemented with 0.1% Tween-20 (ThermoFisher), CD34^+^ iPSC-EP-laden hydrogels were incubated in blocking buffer (DPBS supplemented with 0.1% Tween-20 and 1% BSA) for 30 minutes and then with 1:40 rhodamine-phalloidin (ThermoFisher Scientific) overnight. CD34^+^ iPSC-EP-laden hydrogels were transferred to an μ-Plate Angiogenesis 15-well imaging slide (Ibidi). Z-stacks were acquired at 13μm intervals on a spinning disk confocal microscope (Zeiss Axio Observer Z1 with Yokogawa CSU-X1M). At least 4 regions of interest (∼1.3 mm^2^ cross sections) across the full depth of the hydrogel were acquired per technical replicate and at least two technical replicates were analyzed per experimental condition.

### Length and Connectivity Analysis of Vessel-Like Networks

To analyze the total length, connectivity, and lumen diameter of the networks, we used a computational pipeline previously developed ^12^. In brief, confocal z-stacks are filtered and binarized using ImageJ and then analyzed with a MATLAB script that converts the images into a nodal graph that describes the capillary-like structures as nodes (branch/endpoints) and links (vessels).

### Quantitative reverse transcription–polymerase chain reaction

CD34^+^ hiPSC-EPs were encapsulated in collagen hydrogels at a seeding density of 2.2 million cells/mL. Media was replaced daily with endothelial culture media. One, five, or seven days after seeding, messenger RNA (mRNA) combined from at least 3 gels were isolated with an RNeasy Mini Kit (Qiagen) and then reverse transcribed into complementary DNA (cDNA) with a High-Capacity cDNA Reverse Transcription Kit (ThermoFisher Scientific) according to manufacturer instructions. Quantitative reverse transcription–polymerase chain reaction (qPCR) was performed with PowerUp SYBR green (ThermoFisher Scientific) using the StepOne Plus system (Applied Biosystems). 25 ng of cDNA and 500 nM of primers were used for each reaction. cDNA solutions were activated at 50°C for 2 minutes and then at 95°C for 2 minutes. Forty amplification cycles (95°C for 15 seconds, followed by annealing at 60°C for 60 seconds) were performed. Relative expression of mRNA was quantified using the ΔΔCT method with GAPDH as the endogenous control. Results are presented as mean and standard deviation of three technical replicates. A list of primers used in this experiment is given in Supplementary Table 1.

### Treatment with Viral Spike Protein

Viral spike protein (either SARS-CoV-2 Spike Protein, S1 Subunit, RayBiotech, or MERS-CoV-1 Spike Protein, S1 Subunit, BioVision) was added to endothelial culture media of CD34^+^ iPSC-EP-laden hydrogels at a concentration of 10 μg/mL either one or five days after cell encapsulation. 24 hours after viral spike protein addition, media was replaced with endothelial culture media without spike protein. An overview of this experimental setup is shown in Figure 1. CD34^+^ iPSC-EP-laden hydrogels were cultured for a total of 7 days and then visualized using confocal microscopy (Zeiss Axio Observer Z1 with Yokogawa CSU-X1M).

**Figure 1:**
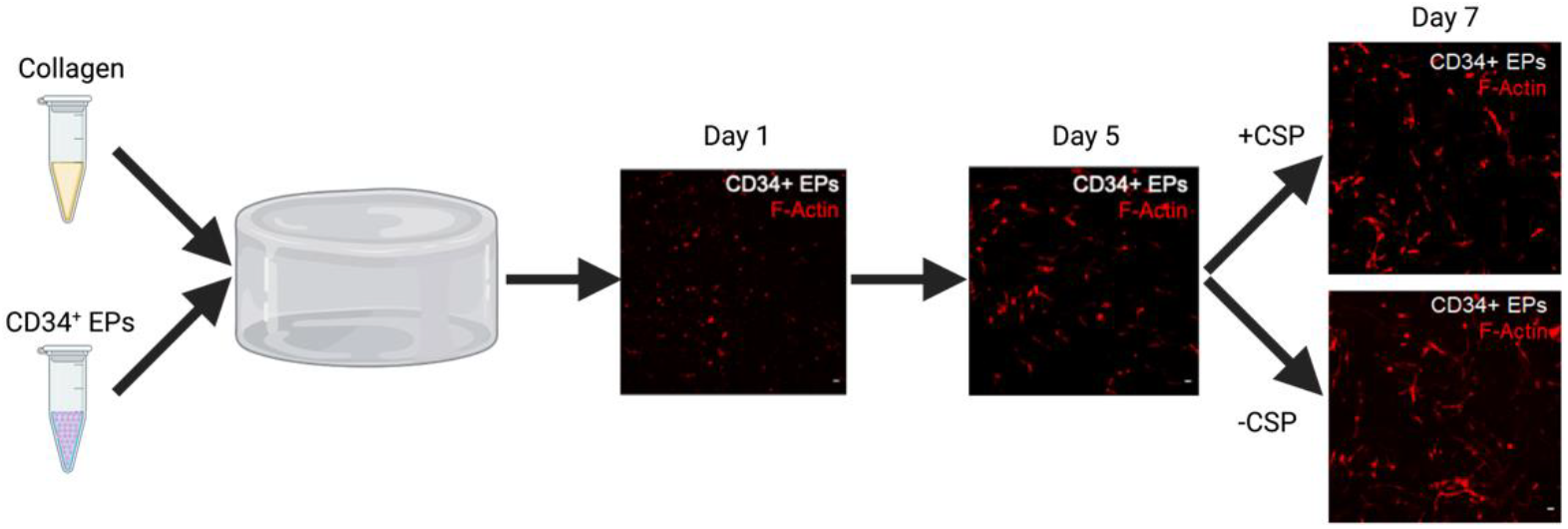
Experimental Overview. CD34^+^-hiPSC-EPs are encapsulated in collagen hydrogels as described previously and cultured for a period of 7 days. During this they extend and form capillary-like networks. 5 days after encapsulation, CD34^+^-hiPSC-EP-laden hydrogels are treated with SARS-CoV-2 Spike Protein (CSP) to simulate COVID-19 infection. CSP treatment results in disruption of vascular networks. Created with Biorender.com

### Dexamethasone treatment

Dexamethasone (MP Biomedicals) was dissolved in dimethyl sulfoxide (DMSO; Sigma-Aldrich) to a working concentration of 25 μg/mL. CD34^+^ hiPSC-EPs encapsulated in collagen hydrogels were cultured for a total of 7 days with daily media changes. 5 days after encapsulation, endothelial culture media was replaced with endothelial culture media supplemented with either 16.6, 25, or 50 ng/mL dexamethasone. 24 hours after dexamethasone addition, media was replaced with endothelial culture media without dexamethasone. 7 days after encapsulation, CD34^+^ hiPSC-EP-laden hydrogels were visualized using confocal microscopy.

### Reducing Endothelial Dysfunction with Dexamethasone

To test if the anti-inflammatory drug dexamethasone would reduce the effects of CSP on CD34^+^ hiPSC-EP-laden hydrogels, 5 days after encapsulation, endothelial culture media was replaced with endothelial culture media supplemented with either 10 μg/mL CSP alone, 16.6 ng/mL dexamethasone alone, or both 10 μg/mL CSP and 16.6 ng/mL dexamethasone. 24 hours after dexamethasone and CSP addition, media was replaced with endothelial culture media without dexamethasone or CSP. CD34^+^ hiPSC-EP-laden hydrogels were cultured for a total of 7 days and then visualized using confocal microscopy. Statistical analysis was performed using two-factor ANOVA followed by a Tukey’s *post hoc* test.

### Cytokine Release Assay

To measure release of inflammatory cytokines following CSP treatment, we used a Proteome Profiler Human Cytokine Array (R&D Systems) using the recommended protocol for cell supernatant. Samples were prepared as followed: CD34^+^ hiPSC-EP-laden hydrogels were cultured for 5 days with daily media changes. 5 days after encapsulation cells were treated with 10 μg/mL CSP for 24 hours. CD34^+^ hiPSC-EP-laden hydrogels not treated with CSP served as a control. Conditioned media from 6 hydrogels per experimental condition was collected after 24 hours and used to run the human cytokine array. The arrays were imaged using a MyECL Chemiluminescence Imager (ThermoFisher Scientific) with a 10-minute exposure time. Relative cytokine levels were quantified using ImageJ.

### Statistical analysis

Unless otherwise stated, all statistical analysis was performed using two-factor ANOVA followed by Dunnet’s test for evaluating multiple comparisons. Significance is denoted as follows: * = p < 0.05, **= p < 0.01, *** = p < 0.005, and **** = p < 0.001. Data are presented as mean ± standard deviation of at least two biological replicates.

## Results

### SARS-CoV-2 Spike Protein Disrupts Vascular Network Formation in CD34^+^-hiPSC-EP-Laden Hydrogels

We first tested how SARS-CoV-2 Spike Protein dosage affects the vascular plexus developed from CD34^+^-hiPSC-EPs encapsulated in collagen hydrogels. 5 days post encapsulation, CD34^+^-hiPSC-EP-laden hydrogels were treated with either 10 or 100 μg/mL SARS-CoV-2 Spike Protein (CSP) for 24 hours. CD34^+^-hiPSC-EP-laden hydrogels not treated with CSP served as a negative control. 10 μg/mL CSP has been used previously to induce endothelial dysfunction, and the 100 μg/mL condition served as an upper limit in case 10 μg/mL was insufficient^14^. Representative images of the vasculature from these conditions (Supplemental Figure 1a-c) demonstrated that in the absence of CSP, a majority of the CD34^+^ cells associate with each other to form a dense capillary-like network. However, following CSP treatment, the vascular network is less dense, and a higher percentage of cells have a round rather than extended morphology. Further analysis from the computational pipeline (Supplemental Figure 1d) shows that a 10 μg/mL CSP treatment resulted in a 22% decrease in the number of vessel end points (p=0.035), a 33% decrease in the number of vessel branch points (p=0.0016), and a 29% decrease in the number of links between vessels (p=0.0059) compared to the control. A further increase in CSP dosage to 100 μg/mL resulted in a 17% reduction in the number of end points (p=0.122), a 24% decrease in the number of branch points (p=0.023), and a 22% decrease in the number of links (p=0.043). Because CSP treatment at both tested concentrations was able to significantly disrupt the vascular networks, a 10 μg/mL CSP concentration was used for all subsequent experiments.

### CSP Toxicity in CD34^+^-hiPSC-EP-Laden Hydrogels is Time Dependent

Next, we wanted to test at what time point during plexus development CSP treatment is most detrimental. CD34^+^-hiPSC-EP-laden hydrogels were exposed to CSP at either one day or five days after cell encapsulation. These two conditions would determine whether CSP prevented networks from forming (day one CSP treatment) or disrupted existing networks (day 5 CSP treatment). For both time points, CSP treatment lasted 24 hours. Confocal microscopy shows a loss of network connectivity in CSP-treated hydrogels (Figure 2b-c) compared to control (non CSP treated) hydrogels (Figure 2a). The results from the computational pipeline show that when CSP is added on day 1, there is a 19% decrease in the number of vessel branch points and a 31% decrease in the volume fraction of the collagen hydrogel containing vessel-forming cells, indicating some toxicity, although neither change is statistically significant (p=0.716 and p=0.333, respectively). However, there is a 77% decrease in the percentage of cells that are part of the largest vessel network, which shows that the CSP has a significant impact on network connectivity (p=0.0011). When CSP is added on day 5, the number of branch points decreases by 59% and the volume fraction decreases by 66% compared to the control, both statistically significant decreases (p=0.0181 and p=0.0074, respectively). The reduction in overall network connectivity is 81%, similar to that of the day 1 treatment. Overall, this indicates that CSP is more toxic when added at day 5, after vessel networks had already started forming.

**Figure 2:**
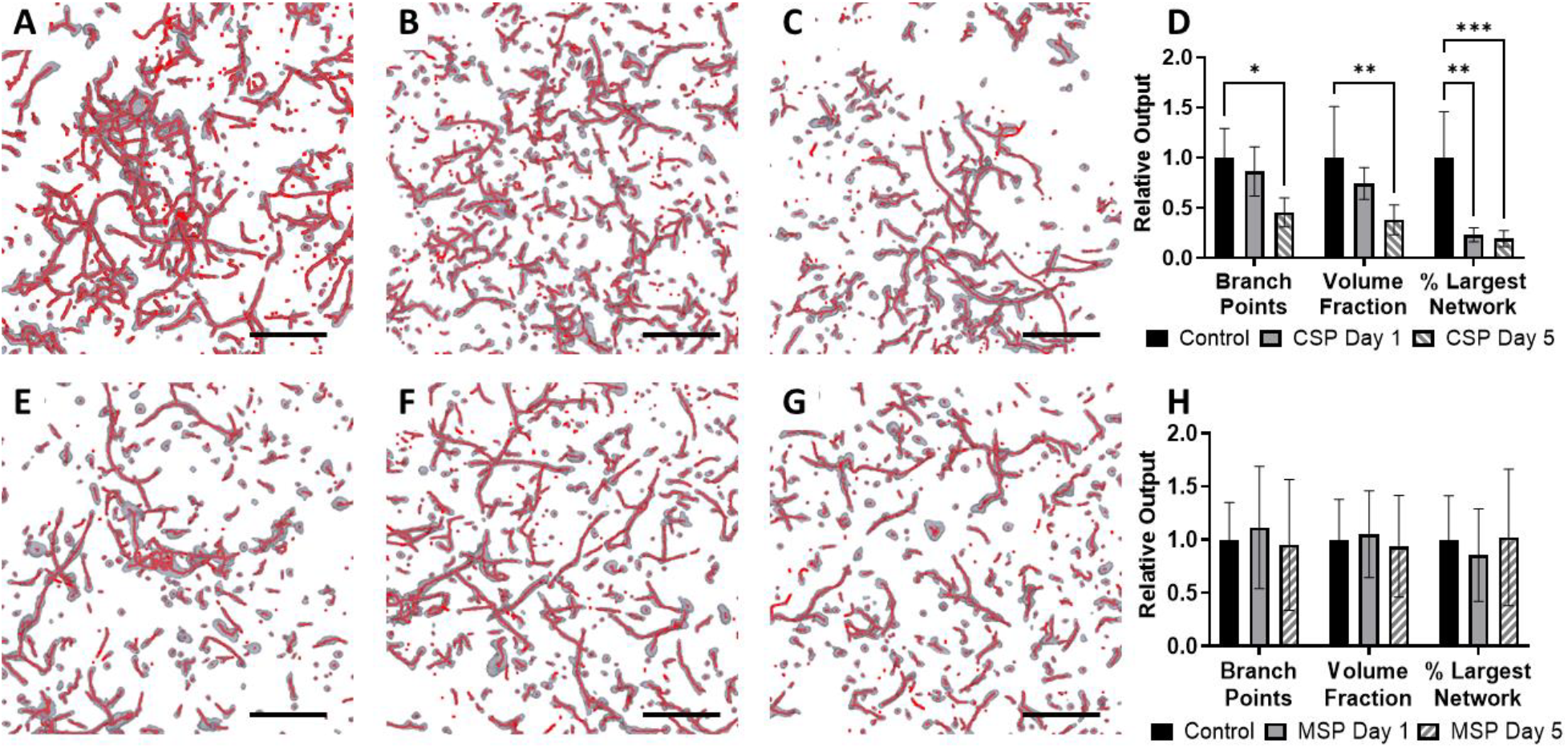
CSP is Toxic to CD34^+^-hiPSC-EPs in Collagen Hydrogels. (a-d) CD34^+^-hiPSC-EPs encapsulated in collagen hydrogels were cultured for 7 days and received either no CSP (a) or CSP for 24 hours 1 day (b) or 5 days (c) after encapsulation. When treated with CSP either on day 1 or day 5, there was a statistically significant decrease in overall network connectivity (% largest network), but there was only a significant decrease in the total number of network-forming CD34^+^-hiPSC-EPs (end points and links) when CSP treatment occurred on day 5 (d). (e-h) CD34^+^-hiPSC-EPs encapsulated in collagen hydrogels were cultured for 7 days and received either no MSP (e) or MSP for 24 hours 1 day (f) or 5 days (g) after encapsulation. There was no statistically significant difference in outputs of the computational pipeline between conditions (h), indicating that the observed toxicity is specific to CSP. Scale bars = 400 μm

### Similar Viral Spike Proteins are not Toxic to CD34^+^-hiPSC-EPs

We next tested whether plexus disruption (as observed in Figures 1 and 2) are specific to SARS-COV-2 or are typical to other coronaviruses treatments, such as the Middle Eastern Respiratory Sickness Coronavirus (MERS-CoV). We chose MERS-CoV because both SARS-CoV-2 and MERS-CoV are able to infect endothelial cells^15,16^ and promote inflammation through toll-like receptor (TLR) signaling^16,17^. However, SARS-CoV-2 also promotes inflammation during viral entry as it binds to ACE2, whereas MERS-CoV does not induce any known signaling following entry into endothelial cells. To test for the effects of TLR-induced endothelial toxicity, we treated CD34^+^-hiPSC-EP-laden hydrogels with 10 μg/mL spike protein from MERS virus (MSP) either 1 or 5 days after encapsulation for 24 hours. We found that MSP treatment did not have any statistically significant effects on vasculature at either tested time point (Figure 2e-h). This suggests that the effects of CSP treatment we observed are specific to SARS-COV-2 virus and not due to TLR signaling.

### CD34^+^-hiPSCs Express ACE2

SARS-CoV-2 can promote inflammation not just through its interactions with TLRs in endothelial cells, but also through SARS-COV-2 binding to its receptor, ACE2^18^. Binding with ACE2 facilitate viral entry and results in downstream signaling that further promotes inflammation. We, therefore, tested if CD34^+^-hiPSCs forming vascular network within collagen hydrogels express ACE2. We followed changes in ACE2 gene expression in CD34^+^-hiPSC-EP-laden hydrogels either 1-, 5-, or 7-days post encapsulation. In addition, we tested multiple endothelial genes (*CD34, CD31, KDR, CDH5)* in order to track endothelial maturation over the 7-day culture period and correlate this to ACE2 expression^10^. RNA isolated from CD34^+^-hiPSC-EPs immediately after sorting served as a control (day 0). *ACE2* expression was at its highest in the day 0 condition. However, only the day 7 condition, which had 62% lower *ACE2* expression compared to the control, was statistically different from day 0 (p=0.0151) (Supplemental Figure 2a). *CD34* is a marker for endothelial progenitors^19^, and we observe a 2.7- and 6.7-fold decrease in *CD34* expression in the day 1 and 5 conditions relative to the day 0 condition (p=0.0005 and p=0.0125, respectively. *CD31* encodes a gene responsible for platelet adhesion to endothelial cells and is used as a marker for mature endothelial cells^20^. *CD31* expression increased by 2-fold, 4.2-fold, and 6.7-fold compared to the control in the day 1, 5, and 7 conditions, respectively (p<0.0001 for all comparisons). Since CD34 is a marker for endothelial progenitors rather than mature endothelial cells, a decrease in *CD34* in combination with an increase in *CD31* indicates that the encapsulated cells are progressing towards mature endothelial cells during the culture period^20^. KDR is the receptor for Vascular Endothelial Growth Factor (VEGF), and its expression remained constant during the 7-day culture period. *CDH5* encodes the protein Vascular Endothelial Cadherin (VE-Cadherin), which is involved in maintaining tight contacts between endothelial cells^20^. *CDH5* expression increased by 2.4- and 2.7-fold compared to the day 0 condition on days 5 and 7, respectively (p<00001 for both comparisons). Other researchers have shown that another coronavirus, SARS-CoV, binding to ACE2 results in ACE2 downregulation^21^. Although SARS-CoV-2 interacts with ACE2 in a similar manner, there is no evidence of ACE2 downregulation. To determine the effects of CSP treatment on ACE2 expression in CD34^+^-hiPSC-EPs, we exposed CD34^+^-hiPSC-EP-laden hydrogels to 10 μg/mL CSP 5 days after encapsulation. 24 hours after CSP treatment, we isolated RNA and performed qPCR with the same genes as described above, with day 0 CD34^+^-hiPSC-EPs as a control. Compared to CD34^+^-hiPSC-EPs not treated with CSP, CSP treatment resulted in a statistically significant increase in *CD31, KDR*, and *CDH5* expression (p<0.0001 for all comparisons). However, CSP treatment did not result in a statistically significant change in *ACE2* expression (p=0.8613) (Supplemental Figure 2b).

### CSP Treatment Induces Release of Inflammatory Cytokines

Previously it has been reported that SARS-CoV-2 binding to endothelial cells results in production of inflammatory cytokines. The inflammatory environment induces apoptosis, which then promotes further cytokine release. This positive feedback loop is known as a cytokine storm^18^. In order to model the cytokines storm in CD34^+^-hiPSC-EP laden hydrogels, we collected conditioned media from CD34^+^-hiPSC-EP laden hydrogels 24 hours after CSP treatment and used a Proteome Profiler Human Cytokine Array to measure relative concentrations of multiple cytokines. We found that CSP treatment resulted in a significant increase in release of interleukin-8 (IL-8, 2-fold over control) and chemokine ligand 1 (CXCL1, 2.9-fold over control), as shown in Figure 3. Both cytokines are produced by endothelial cells^22,23^ and have been shown to be upregulated in COVID-19 patients^24-26^. This data show that CSP treatment creates an inflammatory environment similar to the cytokine storm seen in severe COVID-19.

**Figure 3:**
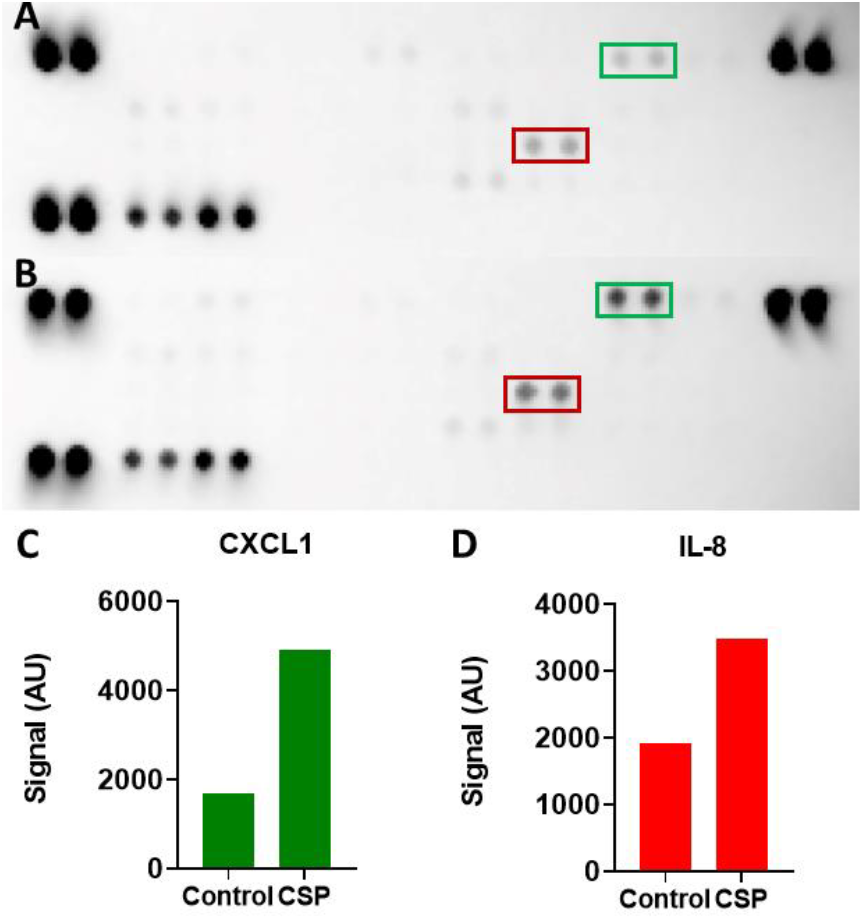
CSP Treatment Induces Inflammatory Cytokine Release. CD34^+^-hiPSC-EP-laden hydrogels were treated cultured for 7 days and either (a) received no treatment or (b) were treated with 10 μg/mL CSP 5 days after encapsulation. Supernatant was collected 24 hours after treatment and concentrations of different inflammatory cytokines were measured using a multiplexed cytokine array. CSP addition resulted in a significant increase in CXCL1 (green/c) and IL-8 (red/d), suggesting that CSP treatment activates endothelial cells.

### Anti-Inflammatory Drug Dexamethasone is not Toxic to CD34^+^-hiPSC-EP-Laden Hydrogels

Dexamethasone is a corticosteroid that has been widely used in clinical settings to treat COVID-19^27,28^. It reduces the effects of the COVID-19 cytokine storm by inhibiting transcription of several inflammatory cytokines. In order to determine if dexamethasone could reduce CSP-induced endothelial toxicity in CD34^+^-hiPSC-EP-laden hydrogels, we first determined a concentration of dexamethasone that was not toxic to these cells. CD34^+^-hiPSC-EP-laden hydrogels were treated with dexamethasone at concentrations of 0, 16.6, 25, and 50 ng/mL 5 days after encapsulation. These are equivalent to the peak serum concentration following a 0, 2, 4, and 8 mg oral dose of dexamethasone, respectively^29^. Computational analysis of the resulting vasculature indicates that the 16.6 ng/mL dexamethasone dosage does not result in a statistically significant change in the vasculature. (Supplemental Figure 3). On the other hand, the 25 ng/mL dexamethasone concentration resulted in a 48.2% decrease in the number of vessel branch points (p=0.0166), and the 50 ng/mL condition resulted in a 45.5% decrease in vessel end points, 67.2% decrease in branch points, and a 53.5% decrease in number of links (p=0.0248, p=0.0005, p=0.0068, respectively). Because the 16.6 ng/mL dexamethasone concentration did not significantly affect vascular network formation, this concentration was used for subsequent experiments.

### Dexamethasone Reduces CSP-Induced Endothelial Dysfunction

After determining a dexamethasone dose that would not affect the vasculature, we then determined if dexamethasone could reduce the toxic effects of CSP. 5 days after encapsulation, CD34^+^-hiPSC-EP-laden hydrogels were treated with either 16.6 ng/mL dexamethasone alone, which serves as a control, 10 μg/mL CSP alone, or a combination of dexamethasone and CSP. Results from the computational pipeline show that CSP treatment results in a 39% decrease in number of branch points, a 36% decrease in the number of links, and a 60% decrease in percent largest network compared to the dexamethasone alone condition (p=0.0352, 0.0497, and 0.0094, respectively), as shown in Figure 4. This shows that CSP is able to both reduce the number of vessel-forming cells as well as vessel connectivity. However, when CSP and dexamethasone are added simultaneously, CSP-induced endothelial dysfunction is not present. In fact, there is no significant difference in any measurement from the computational pipeline between the dexamethasone alone condition and the dexamethasone plus CSP condition. This indicates that dexamethasone is protecting the cells from CSP-induced endothelial dysfunction.

**Figure 4:**
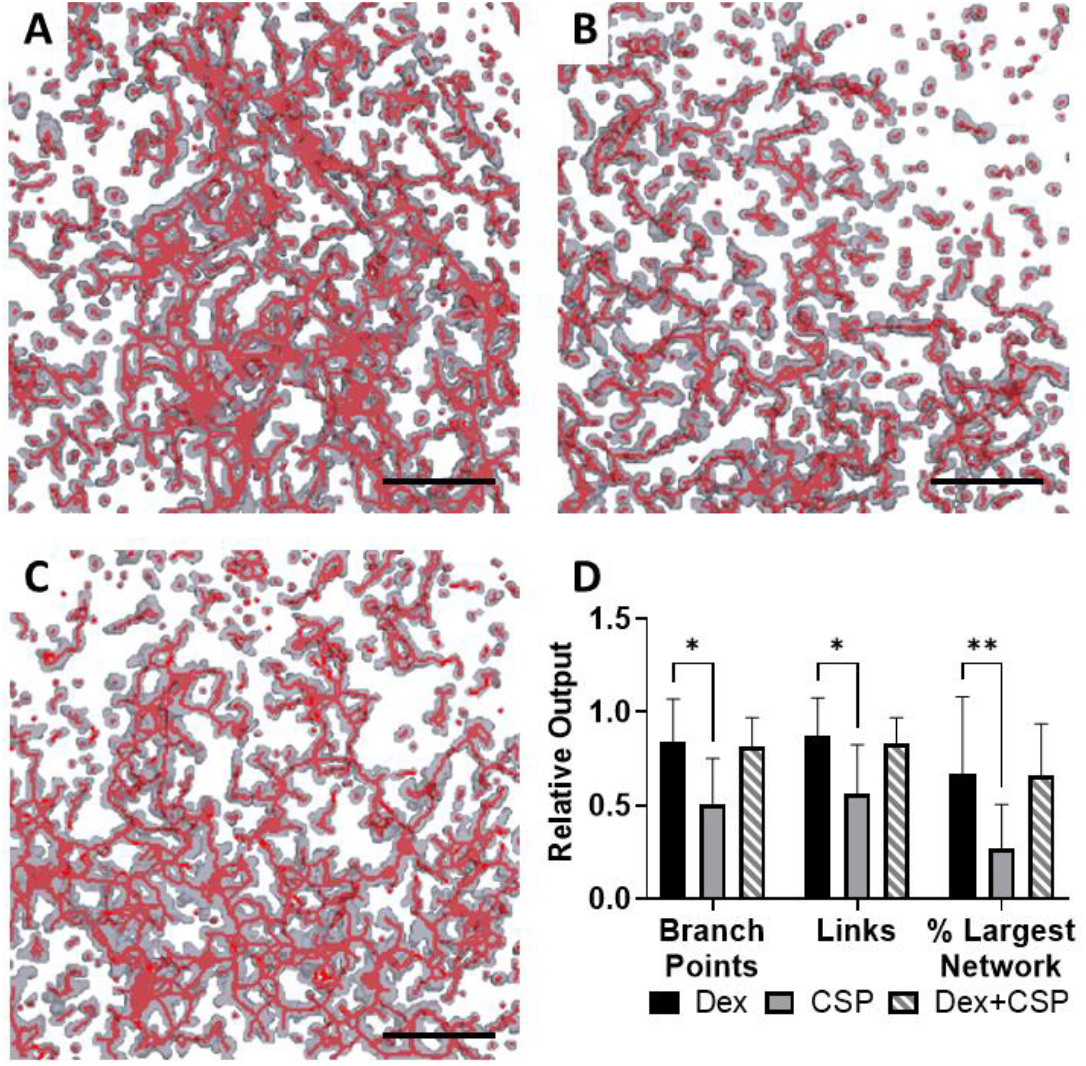
Dexamethasone Reduces CSP-Induced Endothelial Dysfunction. CD34^+^-hiPSC-EPs were encapsulated in collagen hydrogels and cultured for 7 days. 5 days after encapsulation, hydrogels were either treated with 16.6 ng/mL dexamethasone (a), 10 μg/mL CSP (b), or both 16.6 ng/mL dexamethasone and 10 μg/mL CSP (c) for 24 hours. Compared to dexamethasone alone, CSP treatment significantly reduced the number of vessel-forming cells (branch points and links) as well as network connectivity (percent largest network). However, when hydrogels were treated with both CSP and dexamethasone, no outputs from the computational pipeline were significantly different from dexamethasone alone, indicating that dexamethasone can prevent CSP-induced endothelial dysfunction. Scale bars = 400 μm.

## Discussion

The results of this research demonstrate that CD34^+^-hiPSC-EP-laden hydrogels treated with SARS-CoV-2 spike protein can act as a 3D model for COVID-19’s effect on the endothelium. While the effects of COVID-19 on the endothelium have been previously reported, no published research utilizes endothelial cells cultured in 3D or studied the impact of CSP on vasculature. Compared with 2D culture, 3D models produce cells with significant differences in morphology, proliferation, and drug responsiveness^8,9^. Specifically, for COVID-19 modeling, this study shows that CSP can disrupt vascular networks, which is both commonly seen in COVID-19 patients and impossible to replicate in 2D culture.

Most importantly, our model enables study of how CSP affects vasculature formation in a 3D environment. As we previously described, after encapsulation, CD34^+^-hiPSC-EPs extend and then form capillary-like structures over a 7-day period^10-12^; CSP treatment results in the loss of these structures, however the extent of these effects depends on the timing of CSP treatment administration during the process of vasculature formation. Although CSP addition at either day 1 or day 5 resulted in a decrease in network connectivity (percent largest network), CSP addition at a later time point (day 5), once the capillary plexus was formed, results in a significant decrease in the number of vessel-forming cells (branch points and volume fraction). The differences in the number of vessel-forming cells may be due to cell proliferation following CSP removal. The day 1 condition is cultured for 6 additional days after CSP treatment; this allows the CD34^+^-hiPSC-Eps to proliferate to a greater extent than the cells in the day 5 condition, which are only cultured for 2 additional days.

Other researchers have shown that SARS-CoV-2 creates an inflammatory environment both through its interactions with ACE2 and through toll-like receptor signaling^14^. Because TLRs bind a variety of biological molecules other than viral proteins, in order to determine if CSP-induced endothelial dysfunction is due to ACE2 signaling and therefore specific to CSP, we treated CD34^+^-hiPSC-EPs-laden hydrogels with the spike protein from Middle Eastern Respiratory Syndrome (MSP). Although both CSP and MSP associate with TLRs, we did not observe any significant changes in vascular network formation following MSP treatment. This indicates that CSP-induced endothelial dysfunction is due to ACE2 signaling.

In addition to formation of vasculature, CD34^+^-hiPSC-EPs also mature over the 7 days in culture, which is characterized by decreases in *CD34* expression and increases in *CD31* and *CDH5*. This was true for our CD34^+^-hiPSC-EPs at all time points tested. We also measured expression of *ACE2*, the receptor onto which CSP binds for viral entry. We expected *ACE2* RNA to increase with time, as that would provide one possible explanation for the observed increases in CSP toxicity at later time points. However, *ACE2* expression decreased with time relative to day 0 CD34^+^-hiPSC-EPs. In addition, we measured changes in gene expression 24 hours after CSP treatment. ACE2 downregulation requires viral entry, which is primarily mediated by the S2 subunit^30^. Because our experiments only use the S1 subunit, not surprisingly we did not observe any change in *ACE2* following CSP treatment. we would

An alternative explanation is that CSP-induced endothelial dysfunction occurs through an ACE2-independent pathway. Previous research has shown that adult endothelial cells have low expression of ACE2 and that CSP-induced endothelial dysfunction can occur through TLR4 signaling^16^. In addition, CD34^+^ endothelial progenitors have also been shown to express ACE2^31^, so it is possible that the decrease in *ACE2* in the CD34^+^-hiPSC-EPs during the 7-day culture period is a normal part of the cells’ maturation.

In addition to endothelial dysfunction, COVID-19 also results in the release of inflammatory cytokines that results in further endothelial cell death; this is known as a cytokine storm. Some of the major cytokines that play a role in the cytokine storm include IL-1α, IL-1ß, IL-6, and TNFα^24-26^. Although endothelial cells do secrete many cytokines, this is often in response to paracrine effects of immune cells. For example, IL-6 and TNFα secretion by endothelial cells is often associated with IL-1 secretion by immune cells^32,33^. Although this model does not contain immune cells, we observed changes in cytokine secretion from endothelial cells following CSP treatment. Specifically, we found that treating CD34^+^-hiPSC-EPs encapsulated in collagen hydrogels with CSP results in an increase in secretion of IL-8 and CXCL1. Both proteins are secreted by endothelial cells^22,23^ and are present in elevated levels in COVID-19^24-26^. In fact, they both play a similar role in inducing initial migration of neutrophils^34,35^. This suggests that this system can be used to model the early stages of COVID-19 and through coculture with immune cells could be extended to later stages as well.

The corticosteroid dexamethasone is commonly used in clinical settings for treatment of moderate and severe COVID-19 because it reduces the impact of the cytokine storm^27^. Dexamethasone reduces cytokine release by binding to the glucocorticoid receptor, which inhibits transcription of multiple inflammatory cytokines elevated in severe COVID-19, such as IL-1, IL-2, IL-6, IL-8, and TNFα^27^. Because of this, dexamethasone has been widely used in clinical settings for treating moderate and severe COVID-19. We delivered dexamethasone at concentrations ranging from 2 ng/mL to 8 ng/mL to CD34^+^-hiPSC-EPs encapsulated in collagen hydrogels 5 days after encapsulation. In moderate to severe COVID-19, 6-8 mg of dexamethasone is given daily, and the concentrations tested correspond to 25%-100% of the peak serum concentration following an 8 mg dose. We found that a 16.6 ng/mL dose of dexamethasone was the only dose that did not significantly affect vascular network formation. The results of this study were used to test the anti-inflammatory effects of dexamethasone when delivered at the same time as CSP. We treated CD34^+^-hiPSC-EP-laden hydrogels with either 2 ng/mL dexamethasone alone, 10 μg/mL CSP alone, or both dexamethasone and CSP 5 days after encapsulation for 24 hours. We verified that CSP treatment alone resulted in a significant decrease in both number of vessel-forming cells and connectivity. When dexamethasone and CSP were delivered simultaneously, the CSP was not able to induce endothelial dysfunction, and no outputs from the computational pipeline were statistically different from dexamethasone alone. As a result, EPs in collagen hydrogels are able to model the effects of candidate drugs as seen in clinical settings.

In conclusion, we have demonstrated that treating induced pluripotent stem cell-derived endothelial progenitors encapsulated in collagen hydrogels with SARS-CoV-2 spike protein is able to replicate endothelial dysfunction seen in clinical settings. Following 24 hours of CSP exposure, we observed both a decrease in the number of cells forming capillary-like vascular networks as well as a decrease in connectivity of the vasculature. Based on our results this dysfunction is unique to the SARS COV-2 virus and is triggered by release of inflammatory cytokines from the endothelial cells. Same cytokines were measured in patients that had COVID-19 cytokine storm. In addition, we demonstrated that treatment with the corticosteroid dexamethasone is able to block the toxic effects of CSP. These results demonstrate that we can use this system to model COVID-19’s effect on the endothelium to determine the effectiveness of candidate therapeutics or better understand the impacts of long covid in addition, this system can be utilized more broadly for other diseases that affect the endothelium.

### Novelty and Significance

#### What is known?

- Previous clinical data related to the ongoing COVID-19 pandemic has shown that the virus significantly affects the endothelium.
- SARS-CoV-2 binding to the endothelium creates an inflammatory environment that disrupts the endothelial monolayer and results in cell death.
- Most *in vitro* research on SARS-CoV-2 was performed using a monolayer of endothelial cells.

#### What new information does this article contribute?

- We can reproduce in an *in vitro* setting the SARS-CoV-2-induced endothelial dysfunction seen in clinical settings.
- SARS-CoV-2 spike protein promotes inflammation even in the absence of immune cells.
- Treatment with the corticosteroid dexamethasone prevents SARS-CoV-2-induced endothelial dysfunction.

In the ongoing COVID-19 pandemic, researchers have highlighted the virus’s effects on the endothelium. SARS-CoV-2 binding to endothelial cells creates an inflammatory environment resulting in microthrombi, increased vascular permeability, and release of inflammatory cytokines, creating a positive feedback loop known as a cytokine storm. While significant research related to SARS-CoV-2 was performed with endothelial monolayers, these studies do not fully recapitulate the effects on the human vasculature. To the best of our knowledge, we are the first to study the impact of SARS-CoV-2 spike protein on developing 3D vascular plexus. We developed a 3D *in vitro* model of the effects of SARS-CoV-2 spike protein on the endothelium using induced pluripotent stem cell-derived endothelial progenitors encapsulated in collagen hydrogels and treated with SARS-CoV-2 spike protein. We used imaging and an open-source computational pipeline to quantify SARS-CoV-2-induced endothelial dysfunction. In addition, we also observed increases in inflammatory cytokine release following SARS-CoV-2 spike protein treatment and a significant reduction in vascular disruption following treatment with the anti-inflammatory drug dexamethasone. This research highlights the importance of developing *in vitro* 3D vascular plexus models to better understand COVID-19 and Long COVID disease conditions and potentially identify new mitigating therapeutics.

## Acknowledgments

The authors thank Dr. Jeanne Stachowiak (The University of Texas at Austin) for use of the spinning disk confocal microscope, Dr. Sapun Parekh (The University of Texas at Austin) for providing the SARS-CoV-2 spike protein, and Dr. Cody Crosby (Southwestern University) for illuminating discussions on 3D network analysis.

