## Supplemental Figure 1 for "SARS-CoV-2 spike protein induces endothelial dysfunction in 3D engineered vascular networks"

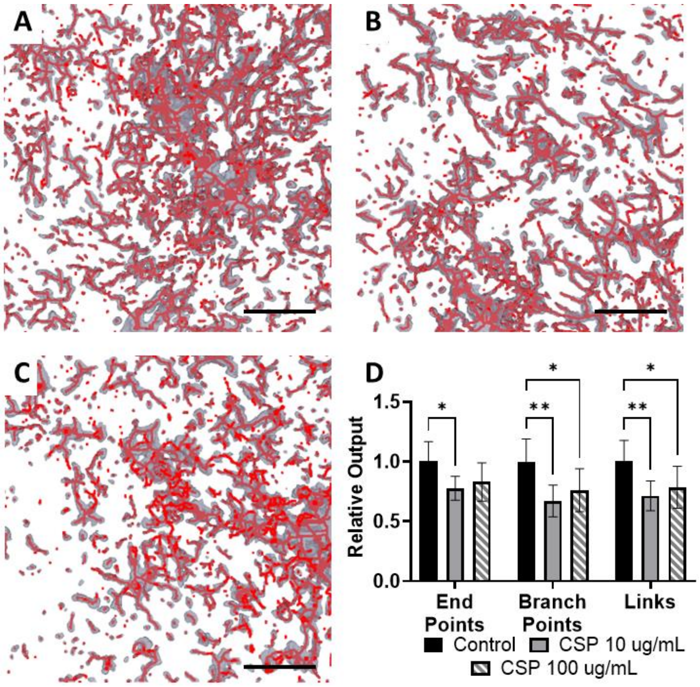


Supplementary Figure 1: Determining CSP Dosage. $CD34^{+}$-hiPSC-EPs were encapsulated in collagen hydrogels and cultured for 7 days. 5 days after encapsulation the hydrogels either received no treatment (a) or were treated with 10 µg/mL CSP (b) or 100 µg/mL CSP (c) for 24 hours. Both the 10 µg/mL and 100 µg/mL conditions had a significant reduction in the number of vessel forming cells, measured by end points, branch points, and links. Scale bars = 400 µm.
