## Supplemental Figure 2 for "SARS-CoV-2 spike protein induces endothelial dysfunction in 3D engineered vascular networks"

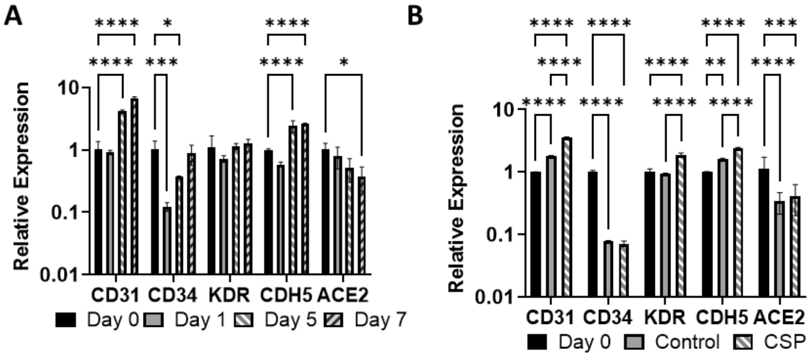


Supplementary Figure 2: Changes in Endothelial Gene Expression. (a) $CD34^+$-hiPSC-EP-laden hydrogel were cultured for up to 7 days. RNA was extracted 1, 5, and 7 days after encapsulation, and RNA extracted from $CD34^+$-EPs immediately after FACS (day 0) served as a control. By the end of the 7-day culture period, there is a significant difference in *CD31* and *CDH5* expression as well as a decrease in *ACE2*. (b) $CD34^+$-hiPSC-EPs were cultured for 5 days and either received no treatment (control) or were treated with 10 µg/mL CSP. RNA was extracted 24 hours after treatment to determine if CSP treatment resulted in ACE2 downregulation, which has been observed by other research groups. We found that the CSP and control conditions had no significant difference in *ACE2* expression, indicating that ACE2 downregulation does not occur following CSP treatment.
