## Supplemental Figure 3 for "SARS-CoV-2 spike protein induces endothelial dysfunction in 3D engineered vascular networks"

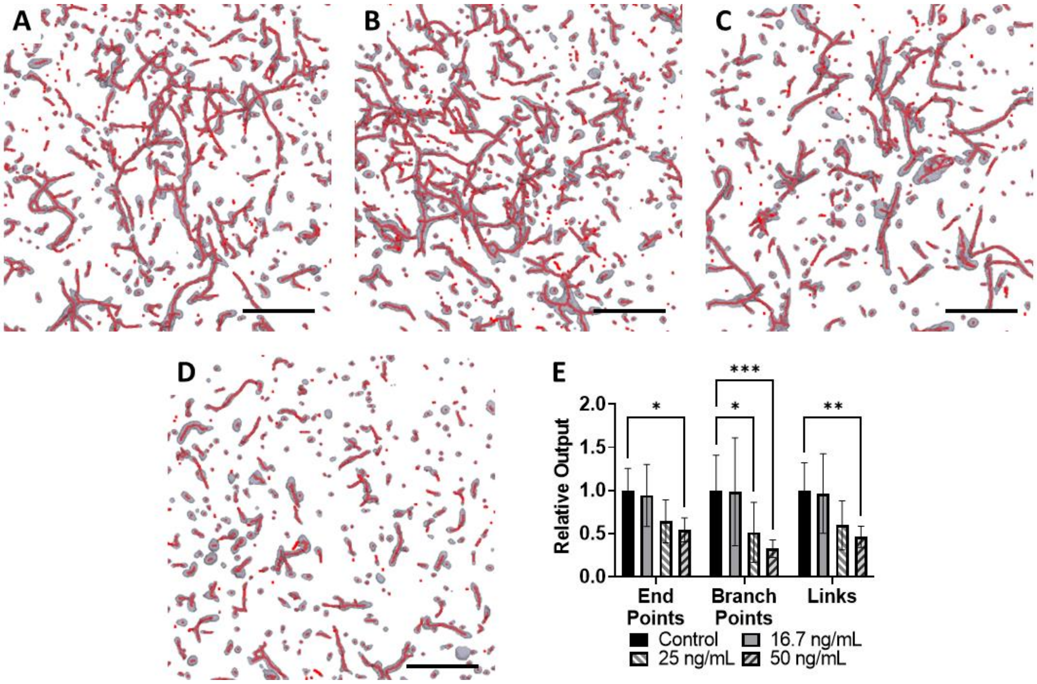


Supplementary Figure 3: Minimizing Dexamethasone Toxicity in iPSC-EP-Laden Hydrogels. iPSC-EP-laden hydrogels were cultured for 7 days and 5 days after encapsulation either received no dexamethasone (a) or dexamethasone at the following concentrations: 16.6 ng/mL (b), 25 ng/mL (c), or 50 ng/mL (d) for a period of 24 hours. The computational pipeline was used to determine a dexamethasone concentration that would not significantly affect the vasculature. There was a significant decrease in one or more pipeline outputs for the 25 ng/mL and 50 ng/mL conditions, but the 16.6 ng/mL condition was not significantly different from the control. Scale bars = 400 µm.
