## Supplemental Table 1 for "SARS-CoV-2 spike protein induces endothelial dysfunction in 3D engineered vascular networks"

Supplementary Table 1: List of qPCR Primers

| Gene | Primer Sequence (F=forward, R=reverse) |
|---|---|
| *ACE2* | Not Available<br>Manufacturer/Catalog Number: Sino Biological/HP100185 |
| *CD31* | F: 5'-GCT GAC CCT TCT GCT CTG TT-3'<br>R: 5'-TGA GAG GTG GTG CTG ACA TC-3' |
| *CD34* | F: 5'-CCT AAG TGA CAT CAA GGC AGA A-3'<br>R: 5'-GCA AGG AGC AGG GAG CAT A-3' |
| *CDH5* | F: 5'-AAG CGT GAG TCG CAA GAA TG-3'<br>R: 5'-TCT CCA GGT TTT CGC CAG TG-3' |
| *GAPDH* | F: 5'-GTC AGT GGT GGA CCT GAC CT-3'<br>R: 5'-CCC TGT TGC TGT AGC CAA AT-3' |
| *KDR* | F: 5'-GTG ACC AAC ATG GAG TCG TG-3'<br>R: 5'-TGC TTC ACA GAA GAC CAT GC-3' |
| *PDGFRB* | F: 5'-TGG CAG AAG AAG CCA CGT T-3'<br>R: 5'-GGC CGT CAG AGC TCA CAG A-3' |
